# HAMAP rules as SPARQL A portable annotation pipeline for genomes and proteomes

**DOI:** 10.1101/615294

**Authors:** Jerven Bolleman, Eduoard de Castro, Delphine Baratin, Sebastien Gehant, Beatrice A. Cuche, Andrea H. Auchincloss, Elisabeth Coudert, Chantal Hulo, Patrick Masson, Ivo Pedruzzi, Catherine Rivoire, Ioannis Xenarios, Nicole Redaschi, Alan Bridge

## Abstract

**Motivation:** Genome and proteome annotation pipelines are generally custom built and therefore not easily reusable by other groups, which leads to duplication of effort, increased costs, and suboptimal results. One cost-effective way to increase the data quality in public databases is to encourage the adoption of annotation standards and technological solutions that enable the sharing of biological knowledge and tools for genome and proteome annotation.

**Results:** We have translated the rules of our HAMAP proteome annotation pipeline to queries in the W3C standard SPARQL 1.1 syntax and applied them with two off-the-shelf SPARQL engines to UniProtKB/Swiss-Prot protein sequences described in RDF format. This approach is applicable to any genome or proteome annotation pipeline and greatly simplifies their reuse.

**Availability:** HAMAP SPARQL rules and documentation are freely available for download from the HAMAP FTP site ftp://ftp.expasy.org/databases/hamap/hamapsparql.tar.gz under a CC-BY-ND 4.0 license. The annotations generated by the rules are under the CC-BY 4.0 license.

**Supplementary information:** Supplementary data are included at the end of this document.

## 1 Introduction

Continuing technological advances have reduced the costs of DNA sequencing enormously in recent years, leading to an explosion in the number of available whole genome and metagenome sequences from all branches of the tree of life [Lewin *et al*., 2018, Mukherjee *et al*., 2017, Paez-Espino *et al*., 2016, Thompson *et al*., 2017, Tighe *et al*., 2017]. This wealth of sequence data presents exciting opportunities for experimental and computational research into the evolution and functional capacities of individual organisms and the communities they form, but fully exploiting this data will require complete and accurate functional annotation of genome sequences. Resources for genome annotation such as RAST/MG-RAST [Meyer *et al*., 2017, Overbeek *et al*., 2014], IMG/M [Chen *et al*., 2017], the NCBI genome annotation pipeline [Haft *et al*., 2018], InterPro [Mitchell *et al*., 2019], TIGRFAMS [Haft *et al*., 2016], and HAMAP [Pedruzzi *et al*., 2015] exploit information from experimentally characterized sequences to infer functions for uncharacterized homologs. While the underlying principles of these resources are undoubtedly very similar, a lack of shared annotation standards and a suitable shared technical framework for annotation hamper efforts to use and combine them.

In this work, we use the HAMAP system (https://hamap.expasy.org) to demonstrate technical solutions that could facilitate the combination and reuse of functional genome annotation systems from any provider. HAMAP classifies and annotates protein sequences using a collection of expert-curated protein family signatures and annotation rules. These rules annotate family members to the same level of detail and quality as expert curated UniProtKB/Swiss-Prot records, combining family membership and residue dependencies to ensure a high degree of specificity. The current implementation of HAMAP uses a custom rule format and annotation engine that are not easy to integrate into external pipelines. The HAMAP-Scan web service (https://hamap.expasy.org/hamapscan.html) is a good alternative for small research projects, but large genome sequencing projects cannot depend on external web services to process large amounts of data. Our goal here was to develop a generic HAMAP rule format and annotation engine that is easily portable by external HAMAP users, using standard technologies that developers of other genome annotation pipelines could also adopt. To do this we have developed a representation of HAMAP annotation rules using the World Wide Web Consortium (W3C) standard SPARQL 1.1 syntax. SPARQL (a recursive acronym for the SPARQL Protocol and RDF Query Language) is a query language for RDF (Resource Description Framework), a core Semantic Web technology from the W3C (see https://www.w3.org/RDF/ for more details). Our implementation allows users to apply HAMAP rules in SPARQL syntax to annotate protein sequences expressed as RDF using off-the-shelf SPARQL engines - without any need for a custom pipeline. If other annotation system providers adopt the same approach, it will be possible to share and combine the annotation rules from different data providers, execute them with any SPARQL engine, and compare the results.

## 2 Methods

To use a generic SPARQL engine to execute rule-based protein sequence annotation, we need the following input data: a) annotation rules in SPARQL syntax, b) protein sequence records in RDF syntax, and c) protein sequence/signature matches in RDF syntax, including alignment information for positional annotations.

To keep the examples given in the Figures short, we provide all RDF namespace prefixes declarations in Figure 1 and omit these from subsequent Figures. We use the UniProt core ontology and other ontologies used by UniProt, such as FALDO [Bolleman *et al*., 2016], which is also used in the RDF of Ensembl [Zerbino *et al*., 2018] and Ensembl Genomes [Kersey *et al*., 2018], to describe sequence positions, and the EDAM ontology [Ison *et al*., 2013] to describe sequence/signature matches.

**Figure 1:**
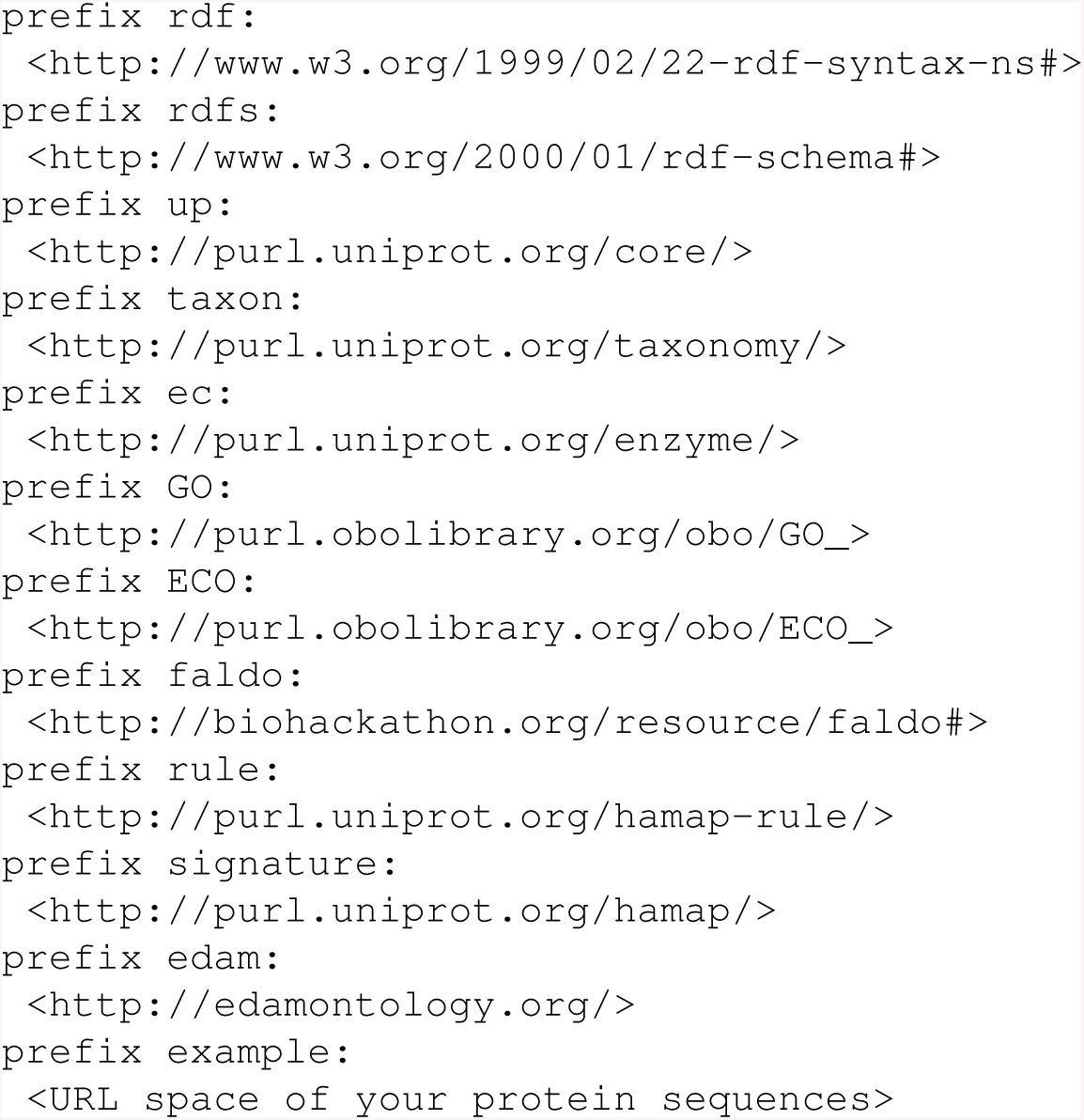
RDF namespace declarations for prefixes used in other Figures.

### 2.1 HAMAP annotation rules in SPARQL syntax

A HAMAP annotation rule consists of two parts: 1) the annotations, and 2) a set of conditions that must be satisfied in order to apply the annotations. The rule annotations can be expressed either by a CONSTRUCT block that returns the annotations as RDF triples or an INSERT block that inserts these triples directly into an RDF store, while the rule conditions can be expressed by the WHERE clause of a SPARQL query. Figure 2 shows part of the HAMAP rule for the signature MF 00005 as a SPARQL query. The CONSTRUCT block generates two RDF triples for two Gene Ontology (GO) terms, providing that all conditions defined in the WHERE clause are satisfied: that the target is a complete protein sequence of bacterial or archaeal origin and is a member of the HAMAP family MF 00005 (i.e. matches the corresponding family signature).

**Figure 2:**
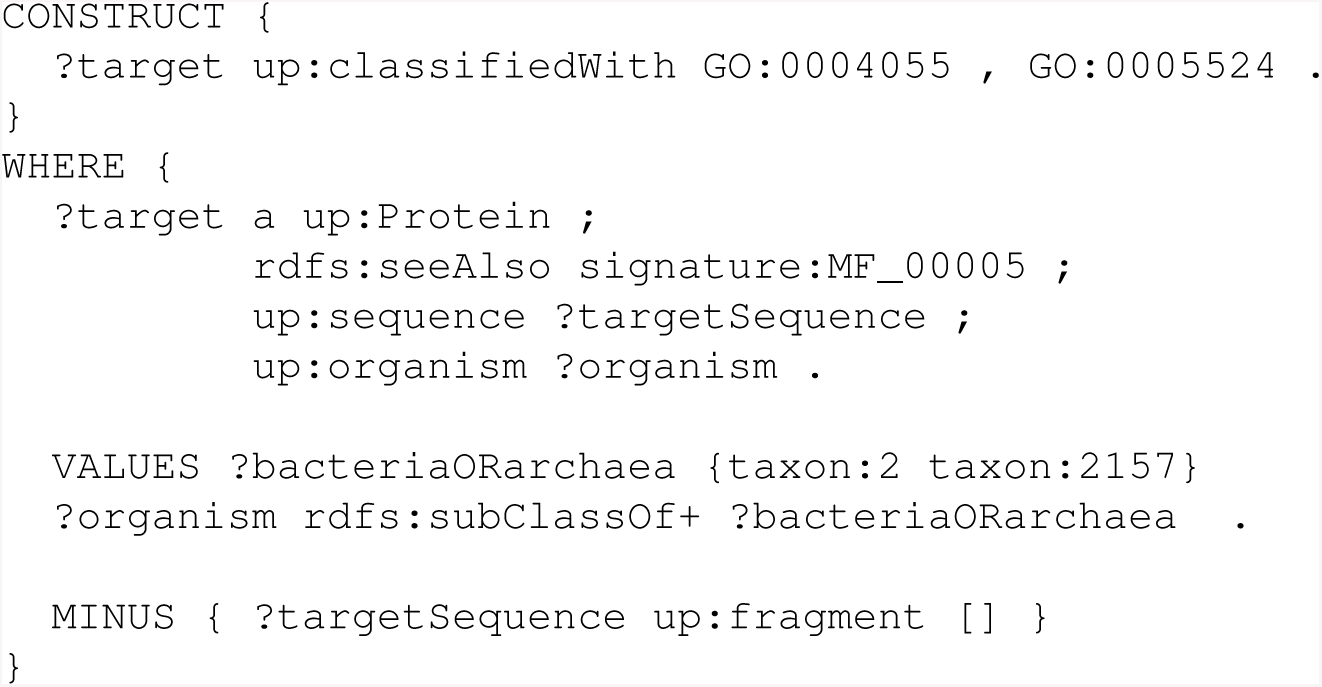
Part of the HAMAP rule for signature MF 00005 as a SPARQL CONSTRUCT query.

Figure 3 shows how the CONSTRUCT block of Figure 2 can be extended to generate metadata for provenance and evidence for each annotation that the rule generates. We attribute the annotations to the HAMAP rule (MF 00005) and describe the type of the evidence with a value from the Evidence Code Ontology (ECO) [Chibucos *et al*., 2016]. We link the attribution to the annotations via RDF reification quads, which is verbose but is understood by all RDF syntaxes and data stores.

**Figure 3:**
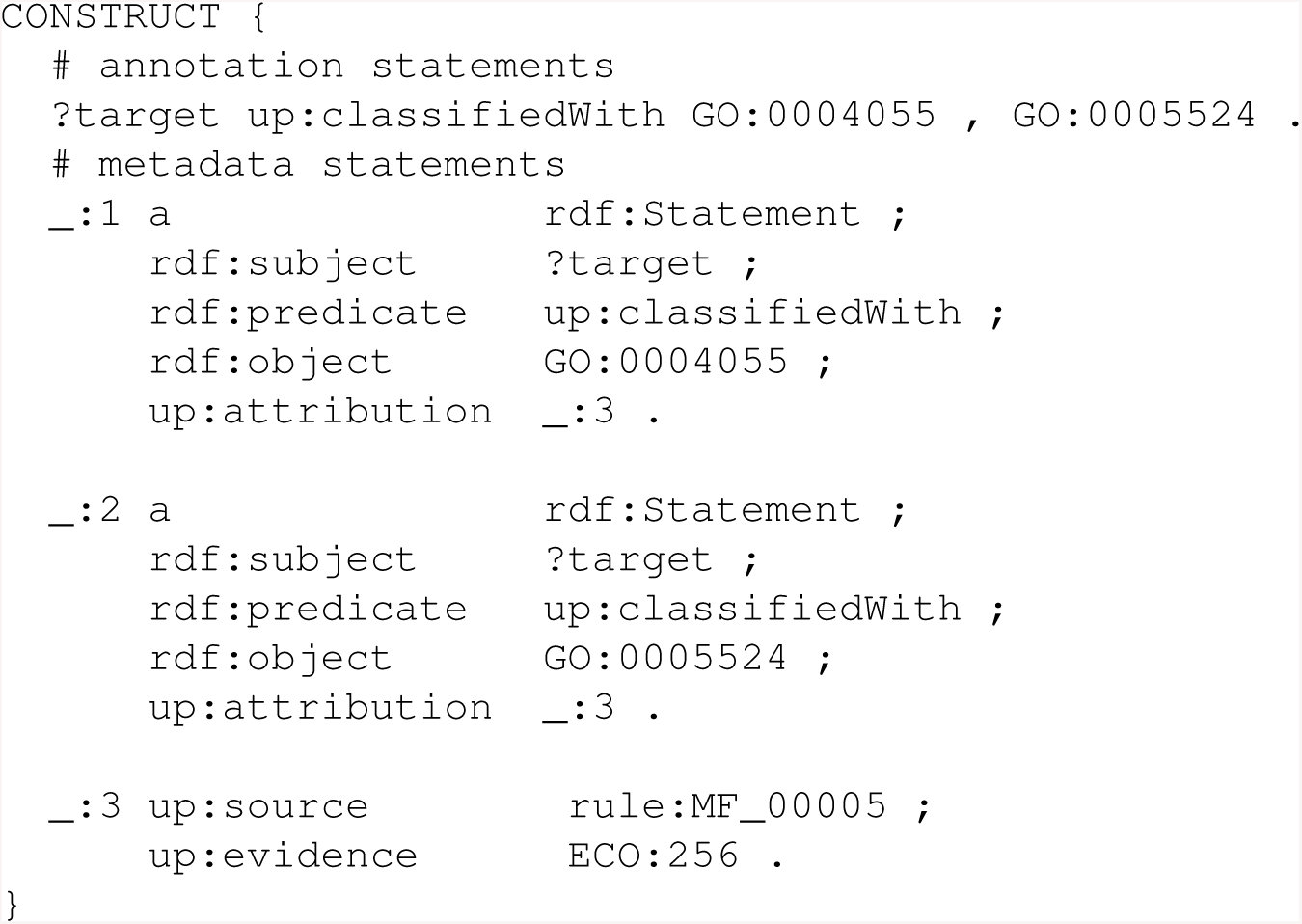
SPARQL CONSTRUCT block of Fig. 2 extended with metadata expressed as RDF reification quads.

The original HAMAP rule implementation has two features that we have not yet implemented in this work. The first is the ability to call sequence analysis methods such as SignalP [Petersen *et al*., 2011] and TMHMM [Sonnhammer *et al*., 1998] for the annotation of signal and transmembrane regions, which is not implemented here as these methods may not be available to external users. The second is precedence relationships between HAMAP rules, which are complex and apply to relatively few rules.

### 2.2 Protein sequence records in RDF syntax

HAMAP SPARQL rules require protein sequence records in a simple RDF format. Figure 4 shows an example protein record with the identifier ‘P1’ (example:P1). The rules require an identifier for the sequence (example:P1-seq) and the organism as an NCBI taxonomy identifier (taxon:83333). The actual protein sequence is provided as an IUPAC amino acid encoded string (in the rdf:value predicate of example:P1-seq) for positional annotations.

**Figure 4:**
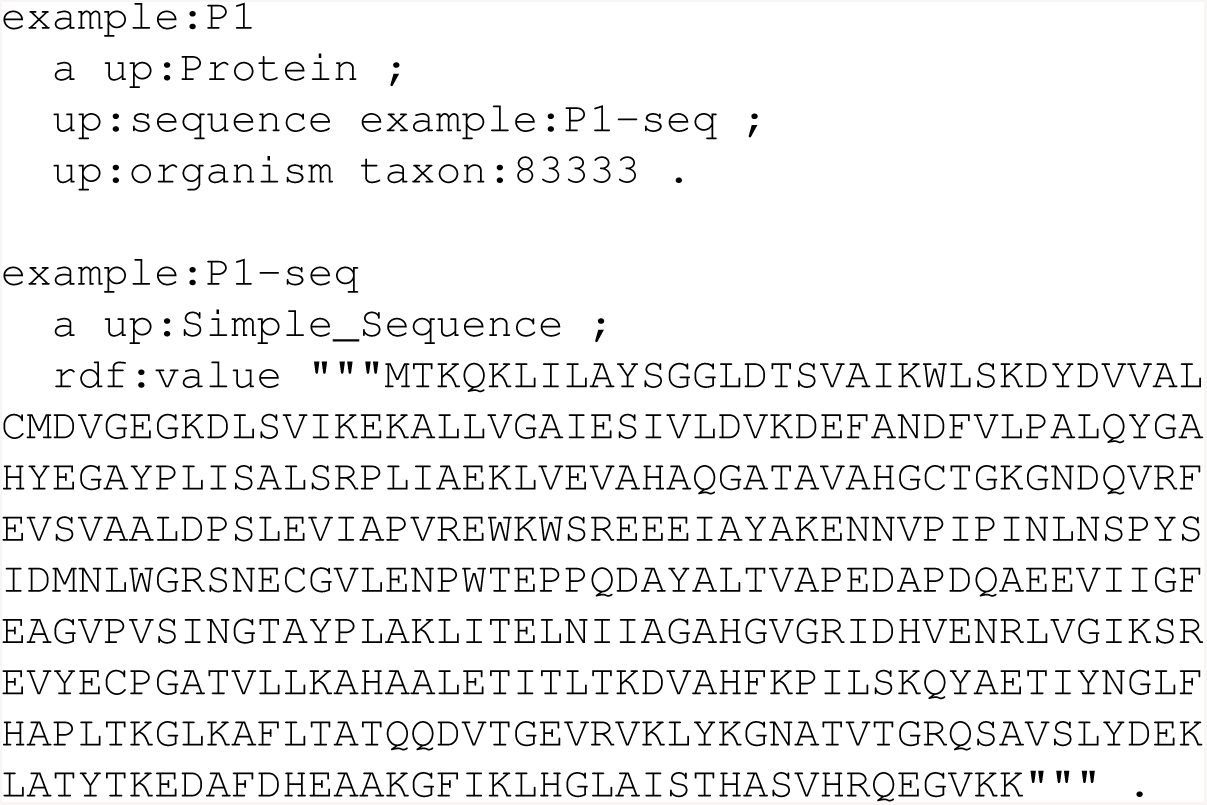
Example protein record in an RDF format suitable for HAMAP SPARQL rules.

### 2.3 Protein sequence/signature matches in RDF syntax

HAMAP SPARQL rules require also sequence/signature match data in an RDF format. Figure 5 shows an RDF representation of the sequence/signature match of the example protein ‘P1’ (Figure 4) and the HAMAP signature MF 00005. The core information is a triple that states that the protein (example:P1) matches the signature (signature:MF 00005). For positional annotations, the rule also needs the start and end positions of the match region on the sequence, as well as the alignment between sequence and signature. We describe this information with the EDAM and FALDO ontologies and use the alignment format returned by the pfsearchV3 [Schuepbach *et al*., 2013] and InterProScan [Mitchell *et al*., 2019] software.

**Figure 5:**
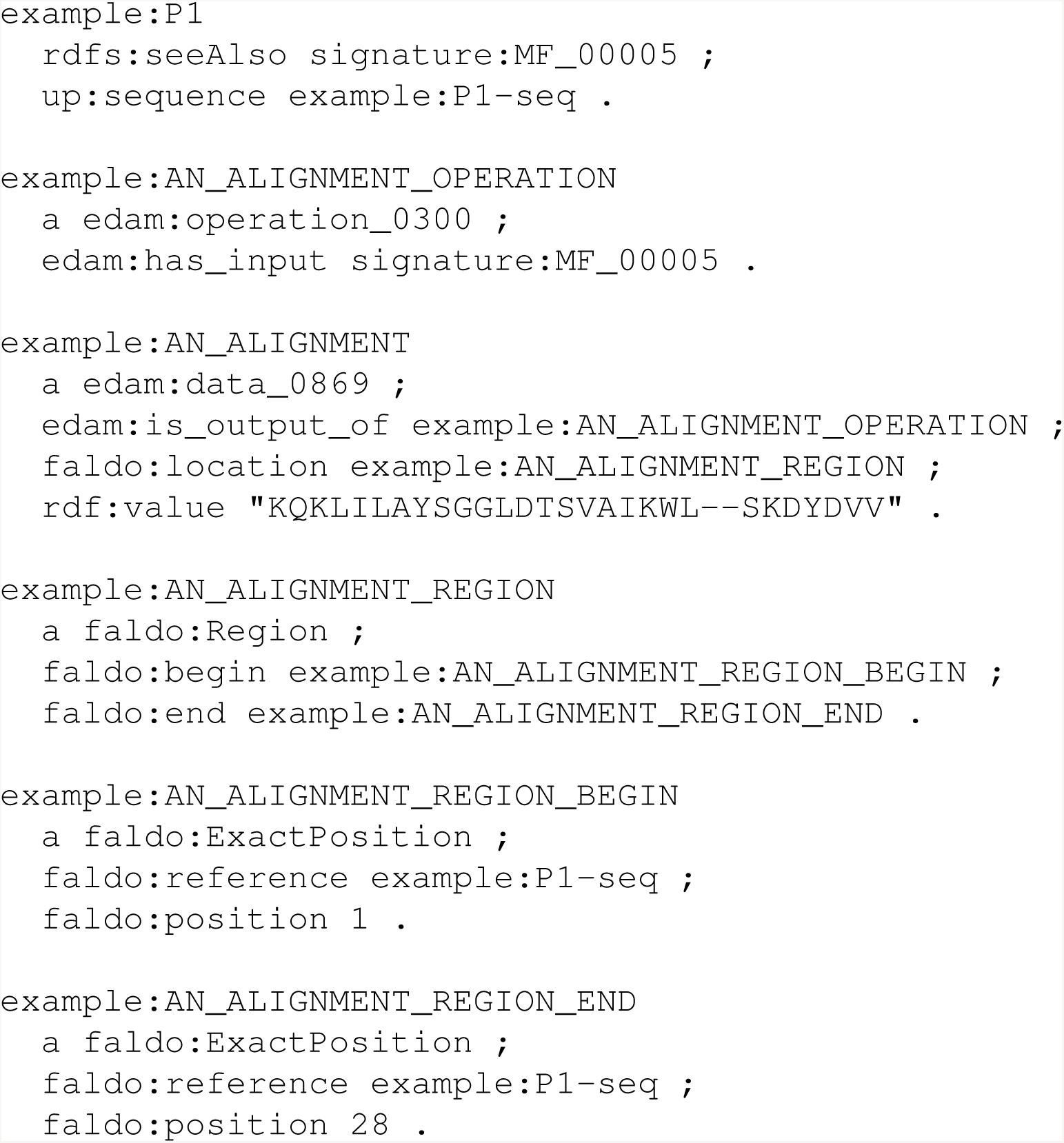
Example protein sequence/signature match in RDF syntax.

HAMAP rules provide a wide range of annotation types, including annotations that apply to specific amino acid positions or ranges on a protein sequence, such as the active site of an enzyme and other functionally important sites and regions. A HAMAP rule specifies the sequence positions of such annotations with respect to one or more experimentally characterized “template” sequences of the HAMAP protein family in UniProtKB/Swiss-Prot. The rule engine therefore needs the alignments of the rule’s signature to its template sequence(s), as well as the alignment of the rule’s signature to the target sequence, to determine corresponding positions on the template(s) and target sequence. A HAMAP rule may additionally require that the position/range on the target sequence matches a specified residue or sequence motif, e.g. to check that an active site has the expected amino acid. This functionality can be implemented either in standard SPARQL 1.1 using the REPLACE, STRLEN and CONCAT functions (see Supplementary Figure 1 for an example), or via a custom SPARQL function (an example Java function for RDF stores that extends the Apache Jena ARQ SPARQL engine is given in Supplementary Figure 2). We distribute the template sequence/signature alignments that are required for rule application together with the rules on our FTP site (at ftp://ftp.expasy.org/databases/hamap/hamapsparql.tar.gz).

### 2.4 Simple rules for standardized annotations

HAMAP rules provide functional annotation in the form of free text and using controlled vocabularies and ontologies developed by UniProt and others. These include the Gene Ontology (GO) [The Gene Ontology Consortium., 2019], the Enzyme Classification of the IUBMB (“EC numbers”) [McDonald *et al*., 2009] represented by the ENZYME database [Bairoch *et al*., 2000], and the Rhea database of biochemical reactions [Lombardot *et al*., 2018] based on the ChEBI ontology [Hastings *et al*., 2016]. For users requiring only a subset of these annotations — such as protein-Rhea links, or protein-GO links — it is possible to translate only the desired annotation types into SPARQL queries. We can also modify the CONSTRUCT/INSERT block of the queries to return the results as simple protein-annotation associations (see Table 1). This tabular result format can easily be loaded into a relational database or JSON-based document store and requires no further investment in a Semantic Web technology stack.

**Table 1:** Simple protein-annotation associations of HAMAP rule MF 00005 for UniProtKB entry B1YJ35.

| Protein | Annotation |
| --- | --- |
| uniprot:B1YJ35 | “GO:0004055” |
| uniprot:B1YJ35 | “GO:0005524” |
| uniprot:B1YJ35 | “GO:0006526” |
| uniprot:B1YJ35 | “GO:0005737” |
| uniprot:B1YJ35 | “ec:6.3.4.5” |
| uniprot:B1YJ35 | “rhea:10932” |

## 3 Results

### 3.1 Validation

We have tested the approach of executing rule-based annotation with a generic SPARQL engine with the data from the HAMAP and UniProtKB/Swiss-Prot releases 2019 02. We translated the 2,304 HAMAP rules into SPARQL CONSTRUCT queries and the 559,228 protein sequences into the RDF format described in Figure 4. We generated the RDF representation of the sequence/signature matches, as illustrated in Figure 5, directly from a relational database containing the results of pfsearchV3 scans of UniProtKB/Swiss-Prot versus HAMAP for our internal HAMAP release pipeline. Other groups could achieve the same result by scanning their protein sequences with InterProScan and converting the XML result files into the described RDF format for sequence/signature matches. We provide an XSLT stylesheet for this conversion in Supplementary Figure 3.

We tested two different open-source SPARQL engines (Virtuoso RDF 7.2 and Apache Jena TDB 3.9.0) to execute our rules and validated the generated annotations by comparing them to those obtained from our custom platform. This platform, implemented in Scala/Java, uses as input files protein entries in FASTA format and HAMAP rules in their custom text format to generate annotations in UniProtKB format (text, XML or RDF). The RDF data generated by the different systems was loaded into separate named graphs of an RDF database for comparisons using SPARQL queries to search for annotations unique to any of the three runs (see example query in Figure 6). The existing custom HAMAP annotation pipeline and each of the two SPARQL engines generated identical annotations, except for those that depend on external sequence analysis methods and the evaluation of HAMAP rule precedence, which we did not implement here as described in section 2.1.

**Figure 6:**
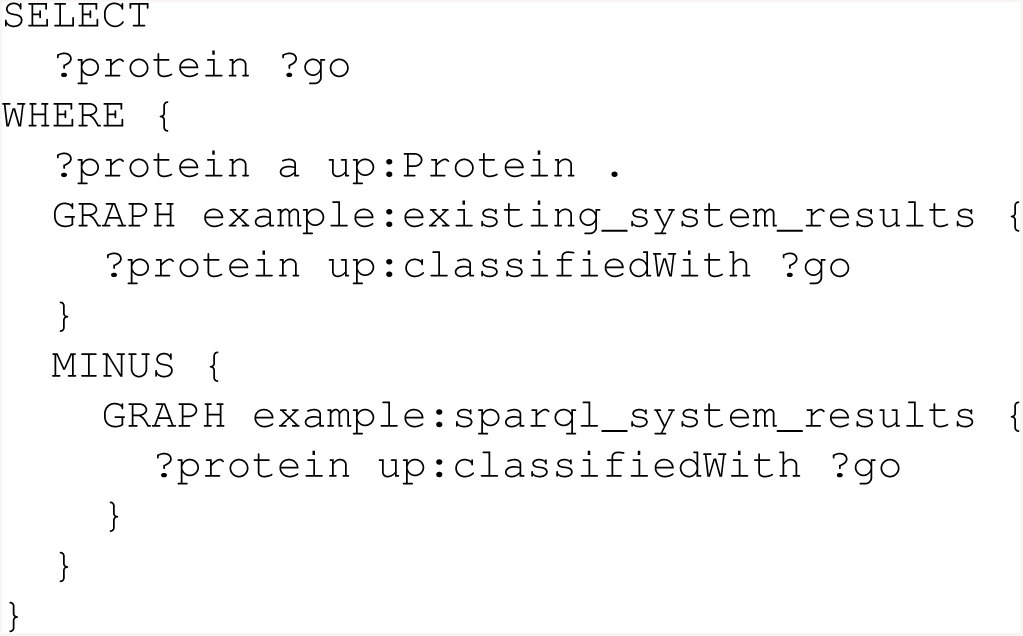
Example query for comparison of GO annotations generated by different systems.

### 3.2 Performance

We executed the SPARQL queries on a 4-year-old desktop with four cores (i5-3470), 16 GB RAM and a single HDD. The annotation of 559,228 UniProtKB/Swiss-Prot sequences, of which 58% match at least one HAMAP signature, took just under six hours. The custom HAMAP platform was significantly faster under these conditions; the setup of an in-memory SPARQL endpoint for each entry in turn dominated the execution time of the SPARQL solution. When running the same queries over all of UniProtKB/Swiss-Prot on a more powerful machine, using a Virtuoso database with all data, we see run times in the seconds for our most complicated rules. In practice, we found no need to optimize the rule application using SPARQL by parallelizing rule execution with more compute nodes, but our approach will benefit from further SPARQL engine optimizations by vendors that develop RDF stores. For now, the application of the rules is faster than the scanning of the sequences with the signatures, which remains the bottleneck in this process.

## 4 Discussion

### 4.1 Protein function annotation pipelines based on SPARQL

Here we have developed a SPARQL representation of HAMAP annotation rules that allows other groups with basic knowledge of this widespread standard technology to apply HAMAP rules in their genome and proteome annotation pipelines. SPARQL can express all features of complex HAMAP rules, including the logic required for positional annotations, while freely available SPARQL engines provide a means to execute HAMAP rules without recourse to specialized software. This work demonstrates the feasibility of adopting SPARQL as a means to integrate existing functional annotation pipelines for genome sequencing projects. This applies not only to expert curated rules from HAMAP and other systems, but also annotation rules generated by automated approaches such as deep learning [Fa *et al*., 2018, Kulmanov *et al*., 2018], which require a feature vector to be expressed as an RDF triple as shown by LOD4ML (http://lod4ml.org). SPARQL can also be adopted by those without access to specialized RDF triple stores by using a SPARQL to SQL mapping (such as that provided by any of the R2RML tools, see https://www.w3.org/TR/r2rml/) to execute SPARQL rules directly against data stored in a relational database. The main weakness of SPARQL is that, like many generic query engines, it tends to be computationally more expensive than a custom solution, but we have seen significant progress in the optimization of SPARQL engines over the past years [Schmidt *et al*., 2010].

### 4.2 An approach that is extensible to any domain of biology

While we have limited our demonstration to the use of SPARQL queries to formalize and execute protein annotation rules from HAMAP, there is nothing that ties the SPARQL approach to a particular domain of biology. Complete genome annotation requires identification and functional annotation of RNAs as well as proteins, and Figure 7 provides a demonstration of how that annotation could be provided by SPARQL. Here a hypothetical SPARQL rule specifies functional (GO) annotation for an RNA sequence of RNAcentral [The RNAcentral Consortium., 2017] that is a member of the U1 spliceosomal RNA family as defined by Rfam [Kalvari *et al*., 2018].

**Figure 7:**
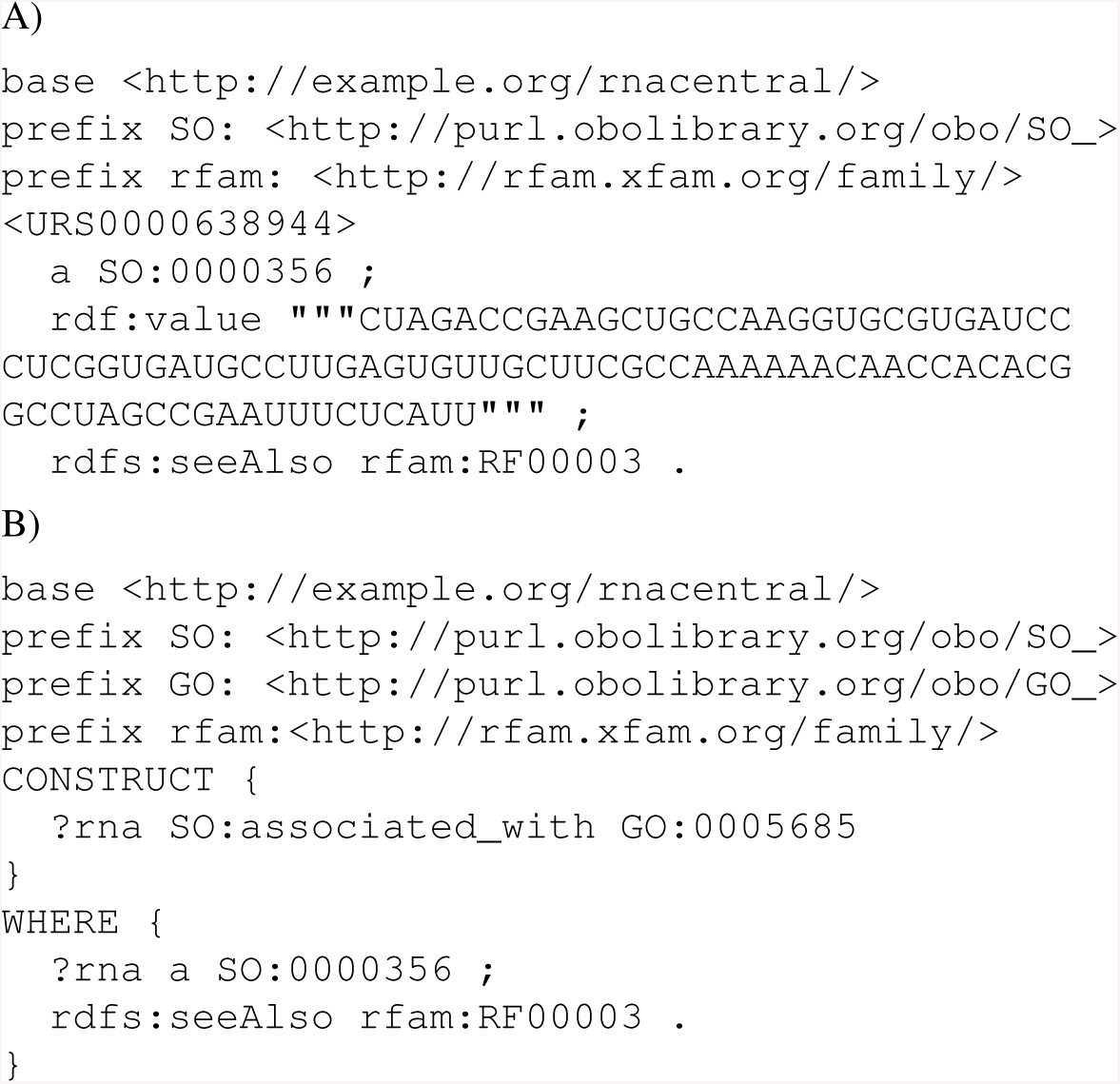
(A) Hypothetical triples to describe a sequence entry from RNAcentral.org that is a member of the Rfam RNA family RF00003 (U1 spliceosomal RNA family). (B) Hypothetical rule associating RF00003 to the GO term GO:0005685 (definition: “A ribonucleoprotein complex that contains small nuclear RNA U1.”).

The development of annotation rules for a given domain across different groups will require community standards for the representation of the relevant domain-specific annotation types. In this work we have used the RDF vocabularies of UniProt, which allowed us to easily compare the results of the SPARQL approach to those of our existing HAMAP rule annotation pipeline. As other appropriate community ontologies become available, our queries and SPARQL rules can be easily adapted.

### 4.3 Further work

We plan to further extend our implementation of HAMAP rules using SPARQL to include external method calls and deal with rule precedence (see Section 2.1), and also develop a SPARQL representation for PROSITE, which provides protein domain annotation via a custom pipeline, ScanProsite (at https://prosite.expasy.org/scanprosite/) [Sigrist *et al*., 2013]. HAMAP and PROSITE are two of the main components of the UniRule system of UniProt, which provides automatic annotation for unreviewed entries of UniProtKB/TrEMBL [The UniProt Consortium., 2019], and the approach described here could feasibly be extended to the entire UniRule system if needed. The UniProt data model was recently extended to allow enzyme annotation using biochemical reaction data from the Rhea database, which will further extend the scope of HAMAP SPARQL rules to more specialized applications that focus on the creation and annotation of draft metabolic models based on reaction networks [Faria *et al*., 2018, Moretti *et al*., 2016].

## 5 Conclusion

## Acknowledgements

We thank Dr. Marco Pagni of the SIB Swiss Institute of Bioinformatics for interesting discussions and critical reading of the manuscript.

## 6 Supplementary information

### 6.1 Supplement 1: Map position on template to target sequence using SPARQL 1.1 standard functions

The InterProScan software represents a sequence/signature alignment in the form of a string where an upper-case letter represents a matched position, a lower-case letter an inserted position and a dash (’-’) a deleted position in the sequence with respect to the signature. This example illustrates how two alignment strings, one for a template/signature and the other for a target/signature alignment, can be used to map a sequence position from a template to a target sequence. This code is not expected to be typed by hand, but generated by tools as needed.

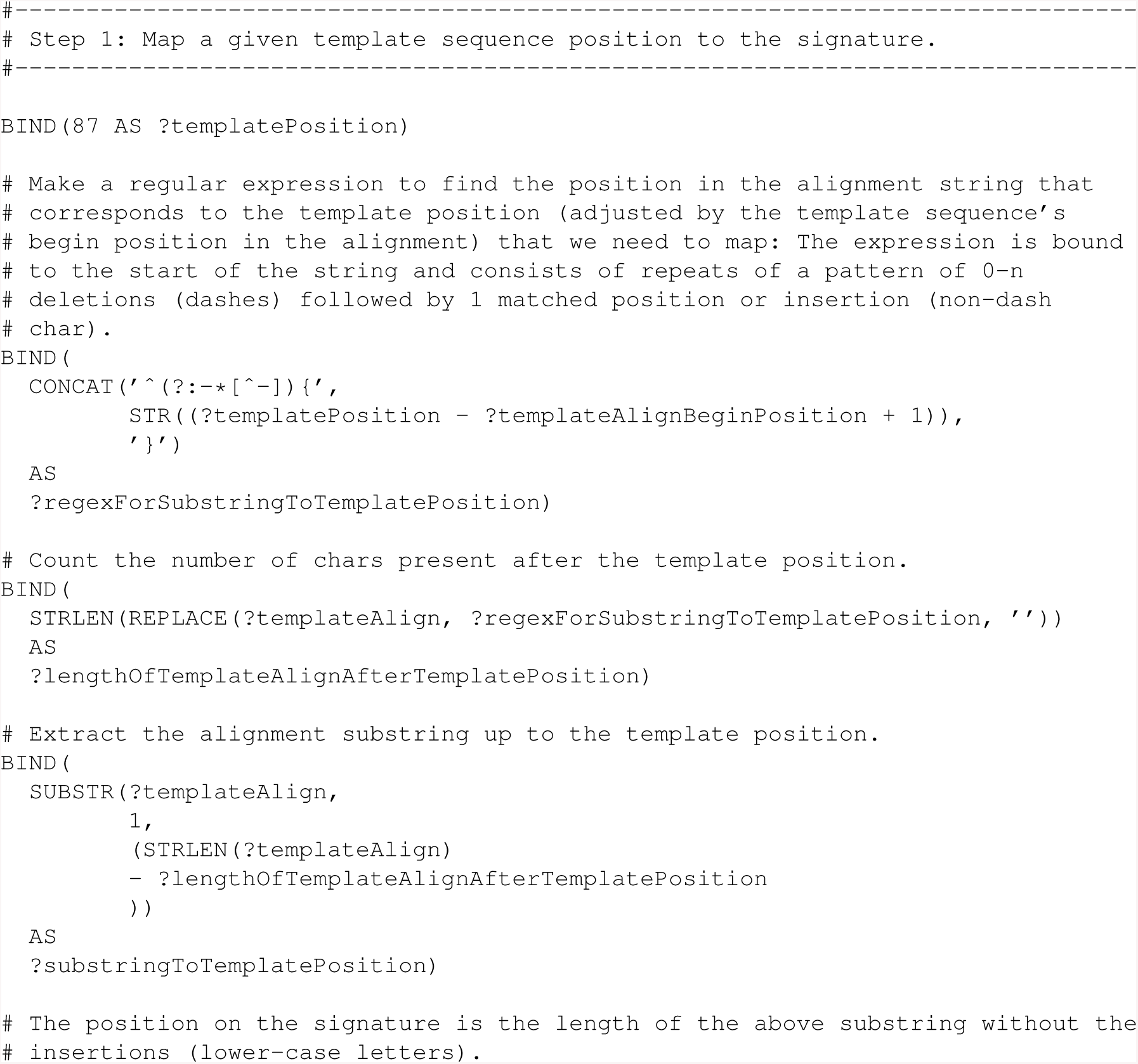
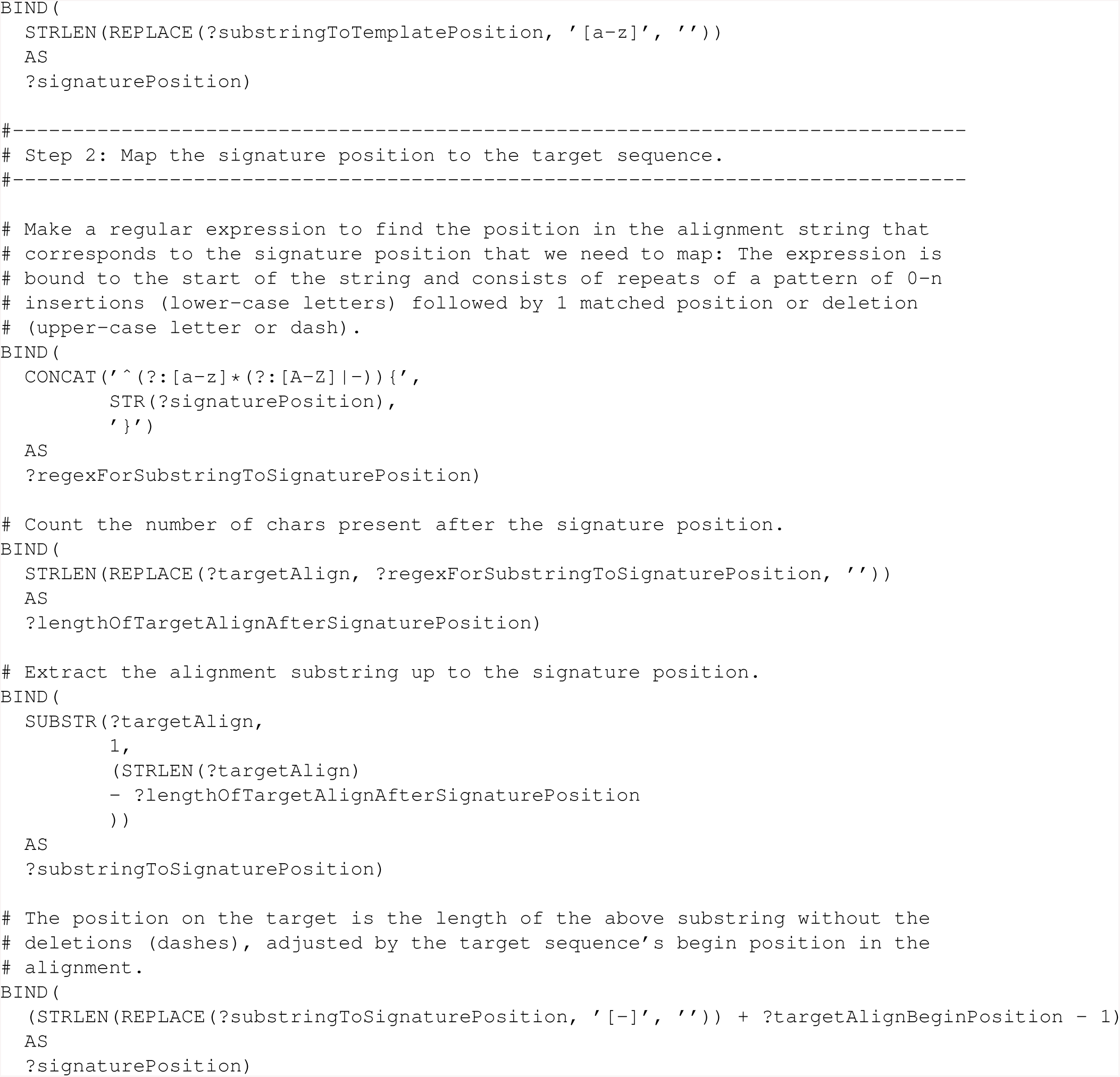

### 6.2 Supplement 2: Java Apache Jena ARQ custom function

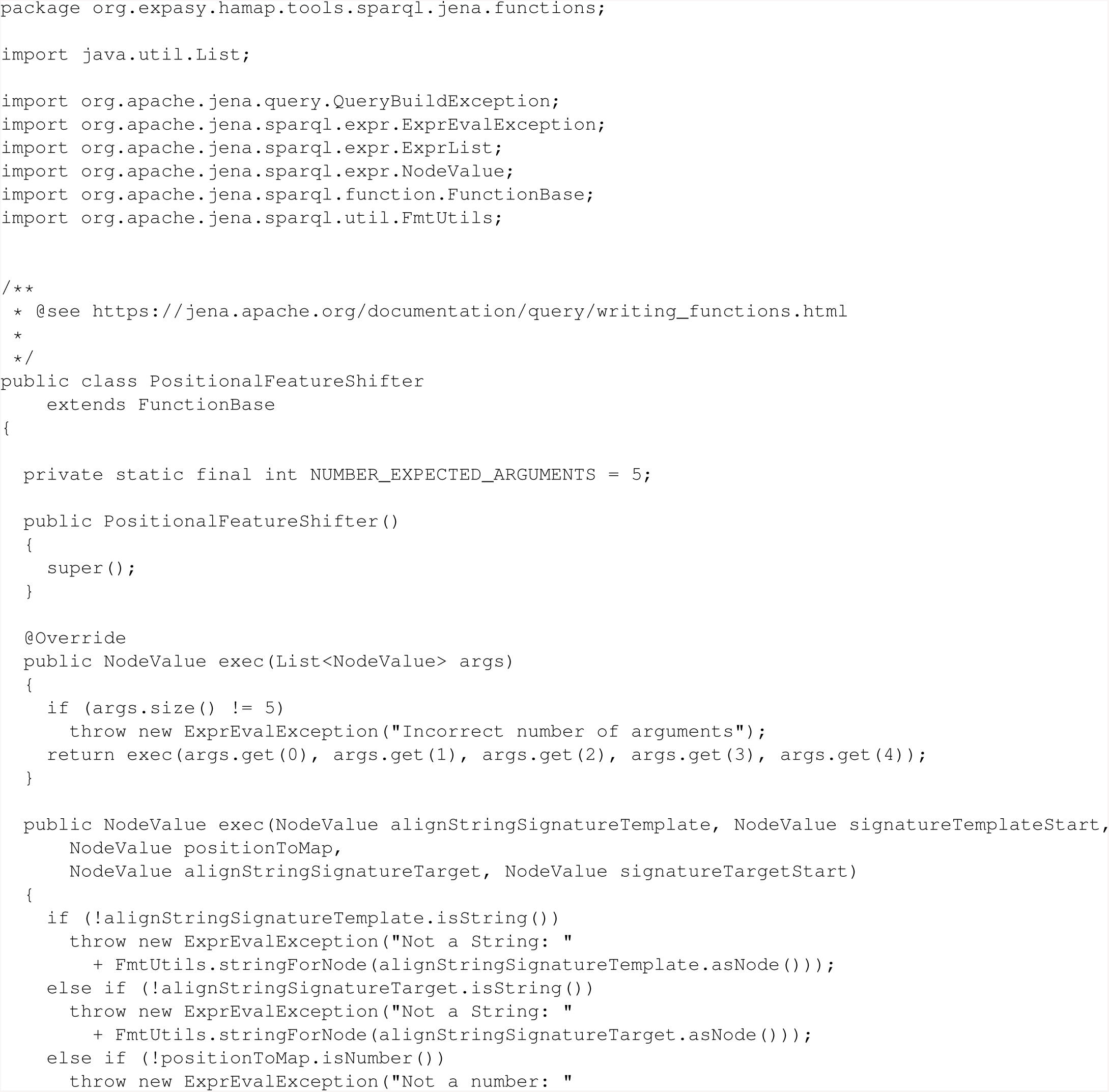
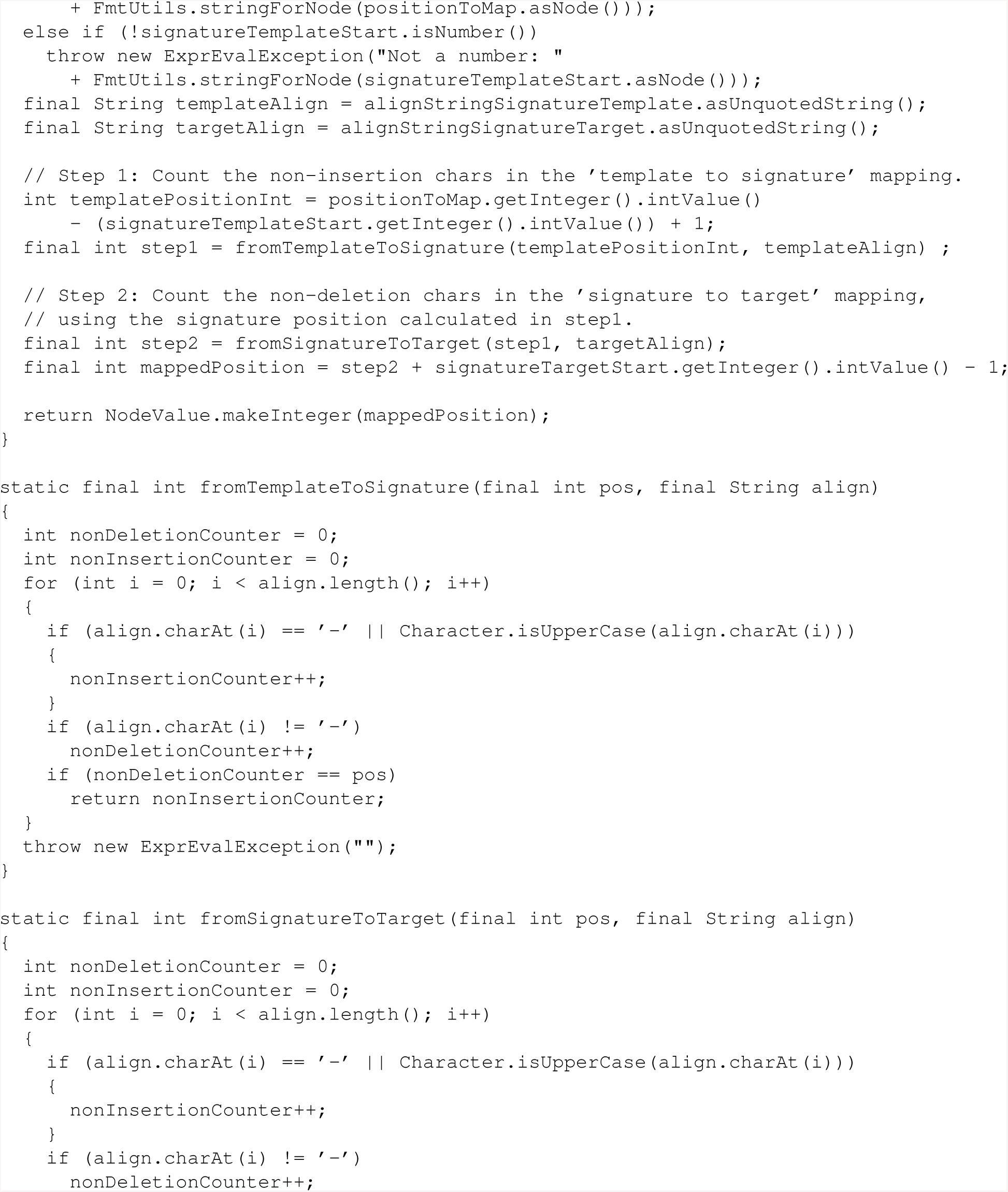
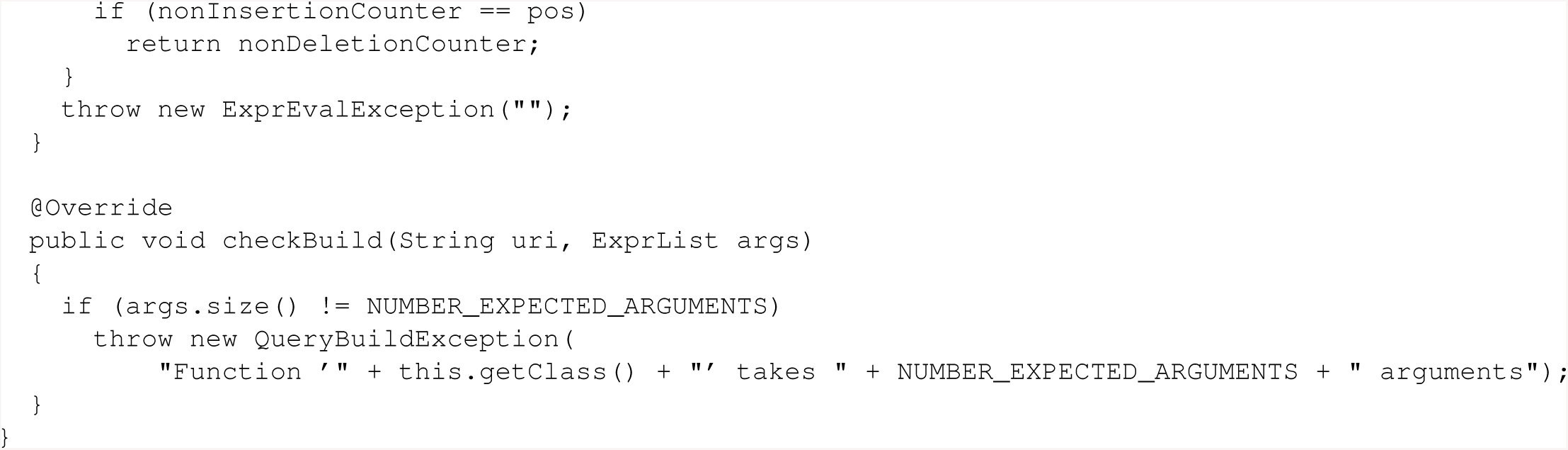

### 6.3 Supplement 3: XSLT to convert InterProScan XML output to minimal RDF for HAMAP

This XSLT stylesheet transforms the XML result file of a local InterProScan run (with the -dp option) into the minimal set of RDF triples required by HAMAP SPARQL rules.

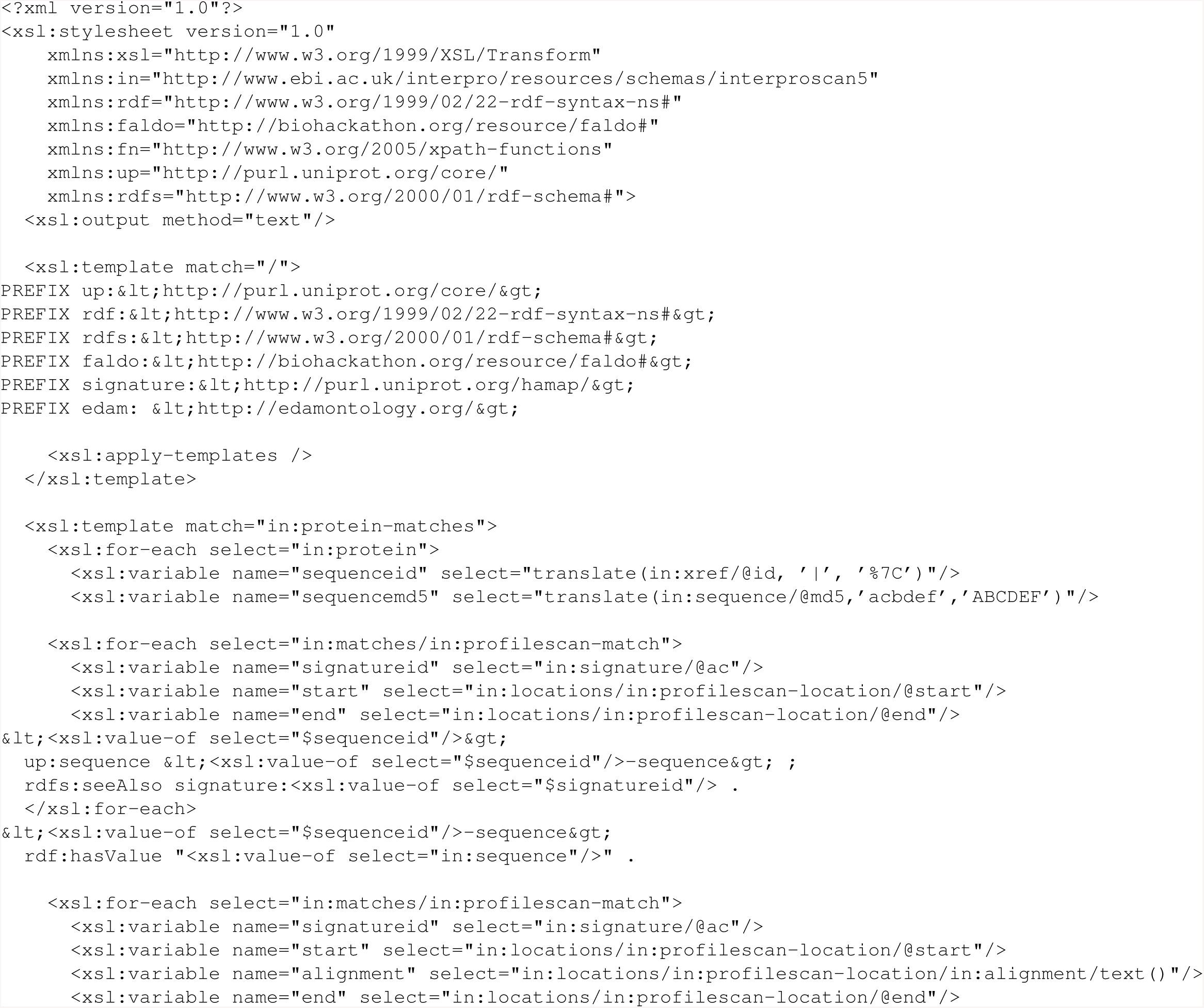
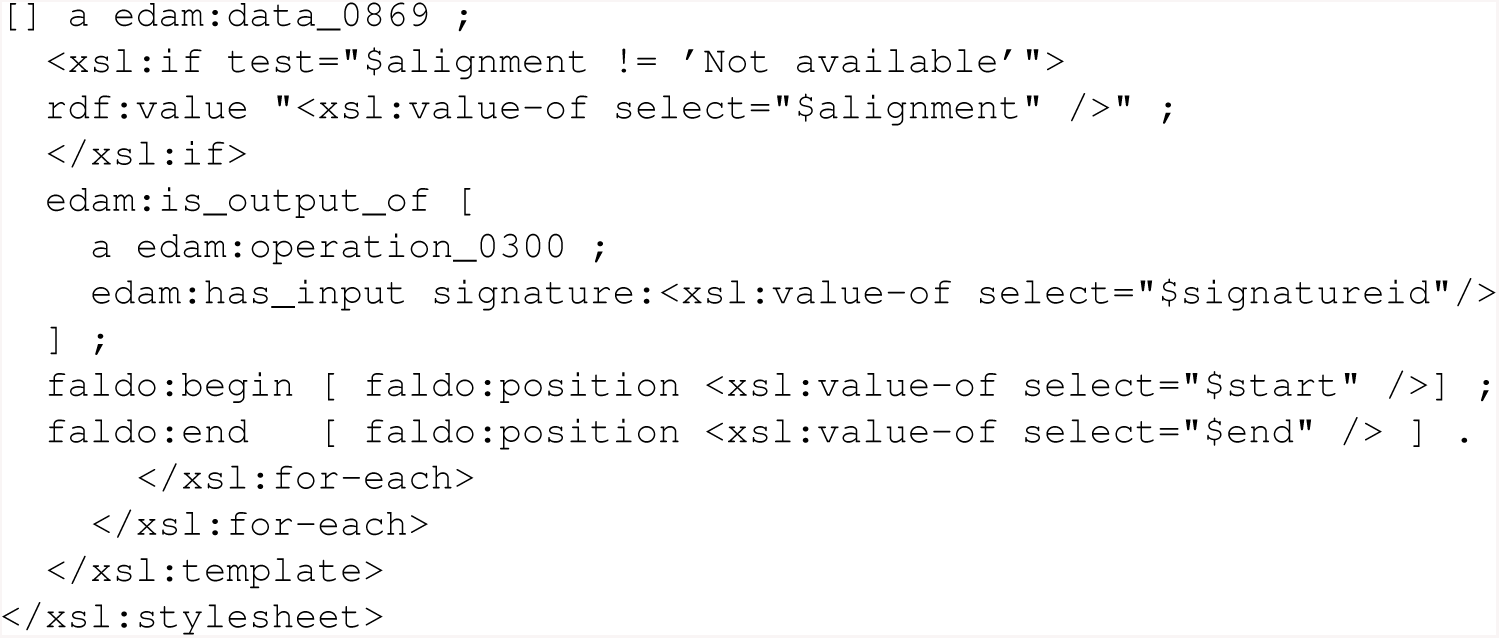

## References

Bairoch, A. The ENZYME database in 2000. Nucleic Acids Res 2000;28(1):304–305.

Bolleman, J.T., et al. FALDO: a semantic standard for describing the location of nucleotide and protein feature annotation. J Biomed Semantics 2016;7:39.

Chen, I.A., et al. IMG/M: integrated genome and metagenome comparative data analysis system. Nucleic Acids Res 2017;45(D1):D507–D516.

Chibucos, M.C., et al. Standardized description of scientific evidence using the Evidence Ontology (ECO). Database (Oxford) 2014;2014.

Haft, D.H., et al. RefSeq: an update on prokaryotic genome annotation and curation. Nucleic Acids Res, Volume 46, Issue D1, 4 January 2018, Pages D851–D860,

Fa, R., et al. Predicting human protein function with multi-task deep neural networks. PLoS One 2018;13(6):e0198216.

Faria, J.P., et al. Methods for automated genome-scale metabolic model reconstruction. Biochem Soc Trans 2018;46(4):931–936.

Haft, D.H., et al. TIGRFAMs and Genome Properties in 2013. Nucleic Acids Res 2013;41(Database issue):D387–395.

Hastings, J., et al. ChEBI in 2016: Improved services and an expanding collection of metabolites. Nucleic Acids Res 2016;44(D1):D1214–1219.

Ison, J., et al. EDAM: an ontology of bioinformatics operations, types of data and identifiers, topics and formats. Bioinformatics 2013;29(10):1325–1332.

Kalvari, I., et al. Rfam 13.0: shifting to a genome-centric resource for non-coding RNA families. Nucleic Acids Res 2018;46(D1):D335–D342.

Kersey, P.J., et al. Ensembl Genomes 2018: an integrated omics infrastructure for non–vertebrate species. Nucleic Acids Res 2018;46(D1):D802–D808.

Kulmanov, M., et al. DeepGO: predicting protein functions from sequence and interactions using a deep ontology-aware classifier. Bioinformatics 2018;34(4):660–668.

Lewin, H.A., et al. Earth BioGenome Project: Sequencing life for the future of life. Proc Natl Acad Sci U S A 2018;115(17):4325–4333.

Lombardot, T., et al. Updates in Rhea: SPARQLing biochemical reaction data. Nucleic Acids Res 2018;47(D1):D596–D600.

McDonald, A.G., Boyce, S. and Tipton, K.F. ExplorEnz: the primary source of the IUBMB enzyme list. Nucleic Acids Res 2009;37(Database issue):D593–597.

Meyer, F., et al. MG-RAST version 4-lessons learned from a decade of low-budget ultra-high-throughput metagenome analysis. Brief Bioinform 2017.

Mitchell, A.L., et al. InterPro in 2019: improving coverage, classification and access to protein sequence annotations. Nucleic Acids Res 2019;47(D1):D351–D360.

Moretti, S., et al. MetaNetX/MNXref–reconciliation of metabolites and biochemical reactions to bring together genome-scale metabolic networks. Nucleic Acids Res 2016;44(D1):D523–526.

Mukherjee, S., et al. 1,003 reference genomes of bacterial and archaeal isolates expand coverage of the tree of life. Nat Biotechnol 2017;35(7):676–683.

Overbeek, R., et al. The SEED and the Rapid Annotation of microbial genomes using Subsystems Technology (RAST). Nucleic Acids Res 2014;42(Database issue):D206–214.

Paez-Espino, D., et al. Uncovering Earth’s virome. Nature 2016;536(7617):425–430.

Pedruzzi, I., et al. HAMAP in 2015: updates to the protein family classification and annotation system. Nucleic Acids Res 2015;43(Database issue):D1064–D1070.

Petersen, T.N., et al. SignalP 4.0: discriminating signal peptides from transmembrane regions. Nat Methods 2011;8(10):785–786.

Schmidt, M., Meier, M. and Lausen, G. Foundations of SPARQL query optimization. Proceedings of the 13th International Conference on Database Theory 2010:4–33.

Schuepbach, T., et al. pfsearchV3: a code acceleration and heuristic to search PROSITE profiles. Bioinformatics 2013;29(9):1215–1217.

Sigrist, C.J., et al. New and continuing developments at PROSITE. Nucleic Acids Res 2013;41(Database issue):D344–347.

Sonnhammer, E.L., von Heijne, G. and Krogh, A. A hidden Markov model for predicting transmembrane helices in protein sequences. Proc Int Conf Intell Syst Mol Biol 1998;6:175–182.

The Gene Ontology Consortium. The Gene Ontology Resource: 20 years and still GOing strong. Nucleic Acids Res 2019;47(D1):D330–D338.

The RNAcentral Consortium. RNAcentral: a comprehensive database of non-coding RNA sequences. Nucleic Acids Res 2017;45(D1):D128–D134.

The UniProt Consortium. UniProt: a worldwide hub of protein knowledge. Nucleic Acids Res 2019;47(D1):D506–D515.

Thompson, L.R., et al. A communal catalogue reveals Earth’s multiscale microbial diversity. Nature 2017;551(7681):457–463.

Tighe, S., et al. Genomic Methods and Microbiological Technologies for Profiling Novel and Extreme Environments for the Extreme Microbiome Project (XMP). J Biomol Tech 2017;28(1):31–39.

Zerbino, D.R., et al. Ensembl 2018. Nucleic Acids Res 2018;46(D1):D754–D761.

